# The Arabidopsis Information Resource in 2024

**DOI:** 10.1101/2023.11.06.565838

**Authors:** Leonore Reiser, Erica Bakker, Sabarinath Subramaniam, Xingguo Chen, Swapnil Sawant, Kartik Khosa, Trilok Prithvi, Tanya Z. Berardini

## Abstract

Since 1999, The Arabidopsis Information Resource (www.arabidopsis.org) has been curating data about the *Arabidopsis thaliana* genome. Its primary focus is integrating experimental gene function information from the peer-reviewed literature and codifying it as controlled vocabulary annotations. Our goal is to produce a ‘gold standard’ functional annotation set that reflects the current state of knowledge about the Arabidopsis genome. At the same time, the resource serves as a nexus for community-based collaborations aimed at improving data quality, access and reuse. For the past decade, our work has been made possible by subscriptions from our global user base. This update covers our ongoing biocuration work, some of our modernization efforts that contribute to the first major infrastructure overhaul since 2011, the introduction of JBrowse2, and the resource’s role in community activities such as organizing the structural reannotation of the genome. For gene function assessment, we used Gene Ontology annotations as a metric to evaluate: (1) what is currently known about Arabidopsis gene function, and (2) the set of ‘unknown’ genes. Currently, 74% of the proteome has been annotated to at least one Gene Ontology term. Of those loci, half have experimental support for at least one of the following aspects: molecular function, biological process, or cellular component. Our work sheds light on the genes for which we have not yet identified any published experimental data and have no functional annotation. Drawing attention to these unknown genes highlights knowledge gaps and potential sources of novel discoveries.

**Article Summary:** The Arabidopsis Information Resource (TAIR, www.arabidopsis.org) is a comprehensive website about *Arabidopsis thaliana*, a small plant that’s very easy to grow and analyze in the laboratory and is used to understand how many other plants function. We share our progress in data collection and organization, website and tool improvement, and our involvement in community projects.

## Introduction

The Arabidopsis Information Resource (TAIR; http://arabidopsis.org) is a comprehensive online digital research resource for the biology of *Arabidopsis thaliana* (Huala *et al*. 2001; Garcia-Hernandez *et al*. 2002; Berardini *et al*. 2015; Reiser *et al*. 2022). The TAIR database contains information about genes, proteins, gene expression, alleles, mutant phenotypes, germplasms, clones, genetic markers, genetic and physical maps, publications, and the research community.

TAIR is a curated database; data are processed by Ph.D.-level plant biologists who ensure their accuracy. Curation adds value to the large-scale genomic data by incorporating information from diverse sources and making accurate associations between related data. Data from manual literature curation, such as protein localization, biochemical function, gene expression, and phenotypes are added to the corpus of knowledge presented for each locus in the genome. TAIR aims to produce a ‘gold standard’ functionally annotated plant genome that plant biologists can use as a reference for understanding gene function in crop species and other plants of importance to humans (Berardini *et al*. 2015). The resource also provides data analysis and visualization tools whose usage has recently been described (Reiser *et al*. 2022).

Initially funded for 14 years by the US National Science Foundation, TAIR has been sustained by subscriptions from academic institutions, corporations, research institutes, and individuals since 2014 (Reiser *et al*. 2016).

For 25 years, a generation of scientists has relied upon TAIR for up-to-date, high quality information and tools provided by scientists and software developers who interact with and respond to the needs of the community. It is a true model organism database used not only by scientists whose primary research organism is *Arabidopsis thaliana* but also by the broader biological research community that uses knowledge gained in this organism to inform their understanding of their organisms. This update covers work done by the resource’s staff in the last few years in the areas of genome functional and structural annotation, tool improvement, FAIR data advocacy, and community service.

## Functional annotation of *A. thaliana* genes using the Gene Ontology (GO)

Since 2001, TAIR curators have been using GO for manual curation of Arabidopsis gene functions from the literature. GO, which describes the biological roles, molecular activities and subcellular localization of gene products, has emerged as the de facto standard for gene functional annotation (Ashburner *et al*. 2000; Gene Ontology Consortium *et al*. 2023). GO curation is the process of extracting and codifying experimental knowledge into annotations that can be used in computational analyses. GO annotations are primarily used to predict functions of unknown genes and newly sequenced genomes, and for gene set analyses for hypothesis generation. TAIR’s ultimate goal is to maintain a gold standard annotated reference plant genome (Berardini *et al*. 2015) that serves as a baseline for predicting gene function in other species, as well as a comparator to other genomes.

GO annotation represents just one aspect of the functional data TAIR curates from the literature. TAIR curators also use the Plant Ontology (PO) to capture gene expression information, craft gene summaries, as well as adding allele and phenotype information, all of which are linked to individual genes. As of October 2023, 13,439 loci have curated summaries, 7,684 loci have one or more phenotypes, 23,123 loci have a total of over 550,000 gene expression annotations to PO terms for gene structure and growth and developmental stages, and 25,500 loci have been linked to primary literature. These counts include information for both sequenced and genetic loci. As genetic loci are cloned, we merge the relevant related records into those of the now known sequenced locus. The locus detail pages in TAIR present a comprehensive view of each locus that includes the data curated from the literature as well as other data sources that help build a more complete picture of an individual locus’ function.

### The importance of curating experimental data

Successful computational methods for inferring gene function invariably rely upon a foundational dataset grounded in experimental evidence. Since many new plant genomes generate their GO annotations based on similarity to Arabidopsis, having a well-annotated genome supported by experimental data is essential to producing high-quality computationally annotated genomes. Each GO annotation includes an evidence code which allows a user to trace whether the supporting evidence is experimental or non-experimental. Annotations with experimental evidence codes are supported by wet lab work, using either low throughput (e.g., BiFC experiments) or high throughput (e.g., proteomics data) methods. Non-experimental annotations are supported by methods that include computational pipelines such as InterPro2GO (Jones *et al*. 2014) that use mappings between domains and functions to assign terms (evidence code of Inferred from Electronic Annotation [IEA]) and phylogeny-based methods like PAINT (Gaudet *et al*. 2011), in which annotations are transferred based on descent from a common ancestor (evidence code of Inferred from Biological aspect of Ancestor [IBA]). In evaluating GO annotations and analysis results that use those annotations, researchers should consider the type of evidence as well as the specificity of the GO term. IEA annotations tend to use more general terms, whereas experimentally supported functions tend to use more specific terms.

### GO annotation datasets for Arabidopsis change over time

GO annotation datasets are subject to change and those changes can affect the analysis and interpretation of data (Jacobson *et al*. 2018; The Gene Ontology Consortium 2019). As with all biological knowledge, what we know about gene function can change over time as new functions are discovered and published. Annotations are also subjected to periodic quality checks to ensure the validity of the data. For example, IEA annotations are removed from the GO after one year and, where possible, replaced with updated (and presumably better) data. All changes to experimentally based annotations are reviewed by curators. Changes to datasets can also occur because of changes to the ontologies themselves (e.g., term inserts, deletes and merges) that necessitate re-examination of the gene-term associations. At TAIR, annotations are updated on a weekly basis on the website and exported on a quarterly basis to the GO where those annotations are merged with Arabidopsis annotations from other sources such as UniProt (The UniProt Consortium *et al*. 2023) and the GO Consortium (GOC) (Gaudet *et al*. 2011). TAIR does integrate the *A. thaliana* annotations made by UniProt and the GOC on a regular basis, as new files are released by these groups. For these reasons, we strongly advise researchers to use the most current annotation data sets either from the TAIR website (https://www.arabidopsis.org/download/index-auto.jsp?dir=%2Fdownload_files%2FGO_and_PO_Annotations%2FGene_Ontology_Annotations) or the GOC website (http://geneontology.org/docs/download-go-annotations/) for any downstream applications such as Gene Set Enrichment Analysis (GESA).

### Current status of gene function annotation in Arabidopsis

We used GO annotation as a proxy to assess the percentage of genes for which some functional information is available. For this analysis, we focused on the proteome (27,657 total, annotation version: Araport11) because the majority of annotations are made to protein-coding genes, and it is the dataset used for orthology-based predictions of gene function.

Before delving into the actual numbers for Arabidopsis, it is important to define what we mean by ‘known’ and ‘unknown’ genes. For each aspect of the GO, we define a ‘known’ gene as one having at least one annotation to that aspect that is supported by an experimental or non-experimental evidence code. ‘Unknown’ genes are defined by having a GO annotation to the ‘root’ term of the ontology, using the ND (No biological Data available) evidence code. For example, a gene product that has no predicted or experimental biological activity would be annotated to the GO term molecular function (GO:0003674), with the evidence code ND, to indicate that the molecular activity of the gene product is unknown at the time of literature review. For each aspect of the GO, we determined (1) the fraction of the proteome that was unknown (UNK), experimentally determined (EXP) and non-experimentally determined (non-EXP) and (2) the numbers and identities of the ‘Unknown’ gene set.

Figure 1 shows a stacked histogram where each bar represents an aspect of the GO. For each aspect, the UNK (yellow), EXP (blue) and non-EXP (red) percentages are shown. The most well annotated aspect is the GO cellular component with 90% of the proteome having either experimental or predicted localization. This is likely due to the relative ease of assaying protein localization and large numbers of proteomics datasets available providing experimental evidence, as well as the relatively facile ability to predict localization based on structural features such as nuclear localization sequences or transmembrane domains. The least well described aspect is GO molecular function, for which 39% of the genome is unknown. Again, this is not surprising considering the difficulty of systematically assessing molecular activities (e.g., a specific enzymatic activity) relative to the generalized biological processes for which those molecular activities are necessary, such as ‘cell proliferation’.

**Figure 1.**
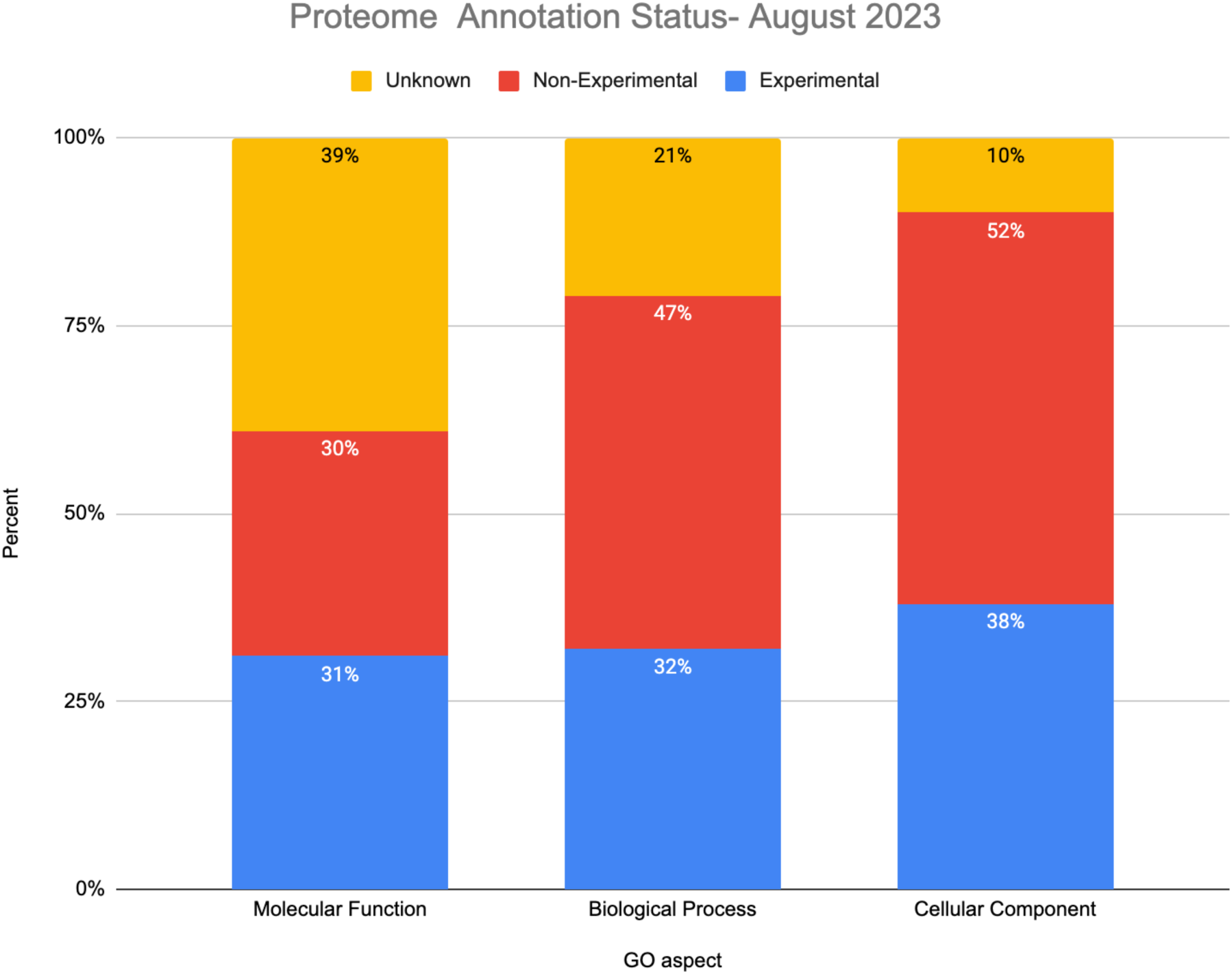
Histogram showing the annotation status of the Arabidopsis proteome by GO aspect and GO evidence class. The unknown set includes proteins with annotations to the root ontology term using the evidence code ND. The experimental set includes proteins with at least one annotation using one of these evidence codes: Inferred from Direct Assay (IDA), Inferred from Expression Pattern (IEP), Inferred from Genetic Interaction (IGI), Inferred from Mutant Phenotype (IMP), Inferred from Physical Interaction (IPI), inferred from High throughput Direct Assay (HDA), inferred from High throughput Expression Pattern (HEP), or inferred from EXPeriment (EXP). The non-experimental set includes proteins ONLY having annotations at least one of the following evidence codes: Inferred from electronic annotation (IEA), Inferred from sequence or structural similarity (ISS), Non-traceable Author Statement (NAS), Traceable Author Statement (TAS), Inferred by Curator (IC), Inferred from Reviewed Computational Analysis (RCA), Inferred from Biological aspect of Ancestor (IBA), Inferred from Sequence Model (ISM).

To identify the set of unknown genes, we sought the intersection of unknowns for each GO aspect. Figure 2 shows a Venn diagram displaying the intersection of unknown genes from each aspect. A total of 1224 protein coding genes lack any functional annotations at all. An additional 3374 lack annotations for biological process and molecular function and thus can also be classified as unknown. 375 of these unknowns are annotated as ‘hypothetical protein’ and may not actually be real genes. Regularly updated versions of this list are available on the TAIR website (https://conf.phoenixbioinformatics.org/pages/viewpage.action?pageId=22807120).

**Figure 2.**
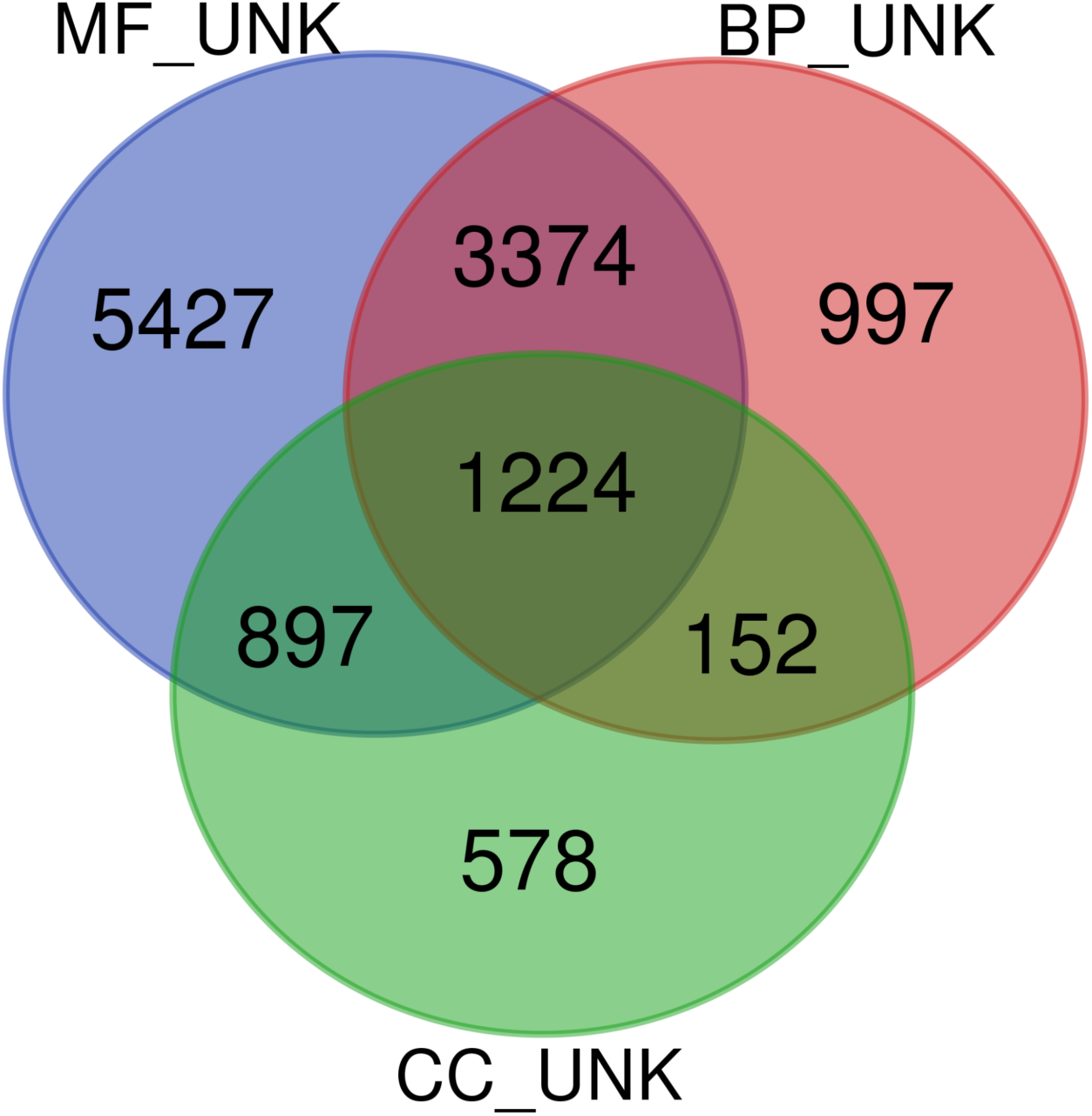
Venn diagram illustrating the overlap among proteins having ND annotations to each aspect. Files containing AGI locus IDs for each aspect (MF_UNK, unknown molecular functions; BP_UNK, unknown biological process; CC_UNK, unknown cellular component; INSERT REF for files) were uploaded to the VIB Venn Diagram Generator (http://bioinformatics.psb.ugent.be/webtools/Venn/).

**Figure 3.**
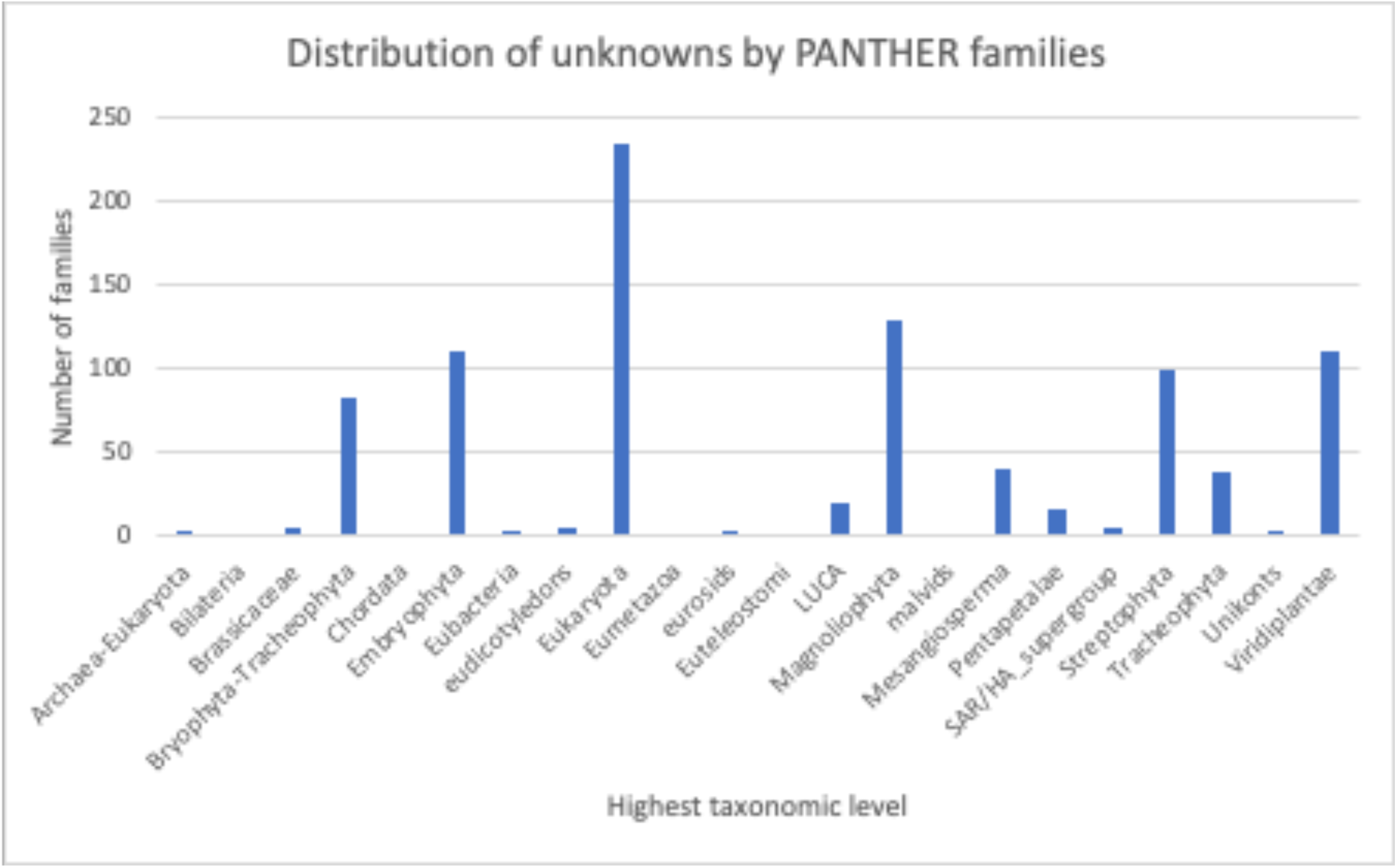
Histogram displaying the distribution of PANTHER 17 gene families containing Arabidopsis unknown proteins grouped by highest taxonomic classification.

Other groups have also used GO as a proxy for known-ness. A different representation of the GO annotation status of Arabidopsis (and other genomes) can be accessed via the Genome Annotation Status Charts (https://genomeannotation.rheelab.org/) generated by Xue and Rhee (Xue and Rhee 2023). Observed differences between the data presented here and in the snapshot may be due to differences in the source files (we used our curation database whereas the Xue paper uses the GAF files from the GOC) and gene set (whole genome vs. proteome only). The smaller set of unknowns presented here is likely because we limited our analysis to protein coding genes.

### Characteristics of the unknown gene set

There are both biological and non-biological reasons why some proteins remain unknown. Among the 4598 ‘unknown’ genes, 375 are annotated as being ‘hypothetical proteins’ in the most recent public annotation version (Araport11) meaning their existence is questionable. A small number of these ‘hypothetical protein’ genes are not present in the in-progress structural reannotation of the *A. thaliana* genome (see community-driven project below) and new genes were added so these numbers will likely be adjusted with the new genome release (The Arabidopsis Col-CC Reannotation Team, in preparation). Biological reasons might include genetic redundancy or difficult to assess phenotypes. Some unknown proteins might belong to members of plant specific gene families and therefore would not have been included in the phylogenetic based inference work done as part of the PAINT project because that focus is on curating families with representatives from the human genome. These plant specific proteins might have novel functions that have yet to be represented within the GO. We used the PhyloGenes resource (Zhang *et al*. 2020) to examine the phylogenetic classifications of the unknown proteins. Among the 4598 proteins, 4070 mapped to 1650 distinct PANTHER families (PantherDB v.17). 70% of those families belong to ‘plant-specific’ families ranging from Viridiplantae to Brassica specific. Most have no associated GO information but some have annotations based on domains. For example, the PTHR34269 family, http://www.phylogenes.org/tree/PTHR34269, has curated members from Arabidopsis and is a plant specific (spanning eudicots) family. This family is characterized by the presence of a B3 domain (Swaminathan *et al*. 2008) and includes well studied members of the ARF/LAV and REM sub families. B3 domain functions in (sequence specific) DNA binding. Therefore it is likely that other unannotated members of the family also have that molecular activity and that annotation would be supported by sequence/phylogenetic analysis. There are many other families such as PTHR10826 (http://www.phylogenes.org/tree/PTHR10826) that span eukaryotes but plant genes (subfamilies) lack annotations even though there are experimental annotations for other non-plant species. This may be because the plant genes are sufficiently diverged into subfamilies and therefore annotations cannot be propagated without supporting evidence from plants. Another reason is that gene functions may have been described experimentally for other plant species in the literature but, because experimental plant GO annotation is limited, that data has not yet been captured as GO annotations. More comprehensive curation of plant gene function would enable propagation of IBA annotations if there was novelty in the plant lineage.

Human and financial resources are other limitations as many functions may be known but have not been captured as a computable GO annotation or what is known is based on prediction and not experimentation. Rocha et al. (Rocha *et al*. 2023), in creating their Unknownome database, assigned a ‘knowness’ score clusters based on a number of factors including GO evidence weights. Again, even well conserved plant specific families may rank low in knownness by their metric. For example, the Dirigent protein family which is found in the Tracheophyta (https://unknome.mrc-lmb.cam.ac.uk/cluster_details/UKP06412/) has well defined experimental functions for some of the member proteins from Arabidopsis and other species (Paniagua *et al*. 2017) but has a standard knownness of 0.0. Probably the reason this cluster comes up with such a low ‘known’ score is because it is a plant specific gene family (restricted to Tracheophyta see http://www.phylogenes.org/tree/PTHR46442). PAINT annotations prioritize PANTHER families with human genes, therefore anything that is plant specific is unlikely to be curated based on biological ancestry.

### How to fill in the knowledge gaps?

Any approach that uses GO annotation as a proxy for knownness has limitations; the most significant being the incomplete nature of GO annotations. The output of experimental data/literature vastly outpaces the ability of curators to process the data both in terms of sheer numbers of genes described and number of articles published. This is especially true for plants where only a fraction of the knowledge published in the literature has been captured. In order to increase the functional annotation coverage of the genome and provide a more comprehensive functional annotation dataset we need to increase throughput through a two-pronged approach that includes (1) curation at the time of publication and, (2) curation of the ‘backlog’ of papers. At TAIR we have been trying to tackle both approaches. To address the first issue we (and others) have developed strategies and tools to encourage authors to curate their data as they publish (Berardini *et al*. 2012; Rutherford *et al*. 2014; Arnaboldi *et al*. 2020; Larkin *et al*. 2021; Reiser *et al*. 2022). We and others are also exploring the use of machine learning and artificial intelligence to assist in data extraction and curation from primary literature (Müller *et al*. 2018; Kishore *et al*. 2020).

### Community curation: Become a GOATherder

Since 2008, we have developed tools that enable researchers to contribute GO and PO annotations to TAIR and expand the gene function knowledgebase beyond what our curators can do. Since then we have processed 12232 annotations from 1692 papers submitted by 1130 community members. From 2013 to 2020 we supported TAIR’s Online Annotation Submission Tool (TOAST) to facilitate community curation of Arabidopsis genes (Li *et al*. 2012). In 2020 we replaced this tool with the Generic Online Annotation Tool (GOAT; https://goat.phoenixbioinformatics.org/). As with TOAST, GOAT is a literature-based curation tool, meaning it is designed for curating experimental gene function data on a per paper basis. Users can contribute annotations for their own or other people’s published works. The GOAT prototype was developed over two years as capstone projects for two cohorts from the Rochester Institute of Technology.

GOAT is a simple web application that allows for basic GO and PO annotations as well as the addition of comments suitable for incorporation into gene summaries (Fig. 4). GOAT uses ORCiD authentication (https://orcid.org/) so users must register or have an ORCID ID to begin. Once logged in users enter the DOI or PubMed ID for the article they wish to curate. Then they can add as many genes as they want using one of the allowed name times (AGI Locus ID, UniProt ID or RNA central ID). To annotate a gene they must first select the ‘subject’ gene product (from the supplied list) and then a type of annotation (GO biological process, GO molecular function, GO cellular component, PO structure, PO developmental stage, protein-protein interaction or Comment). Based on that selection users can then search for GO or PO terms within that subset of the ontology. They can then pick an appropriate evidence code from the ECO (Evidence and Conclusion Ontology; (Nadendla *et al*. 2022)) ontology for the experiment that supports the assertion/annotation. The interface is intuitive and a tutorial is available on TAIR’s YouTube Channel (https://www.youtube.com/watch?v=t5oB51yX6Lobrief). Community annotations are reviewed by a TAIR curator to make sure that they are consistent with annotation rules and best practices before integration into TAIR and eventual consumption by GOC and other resources.

**Figure 4.**
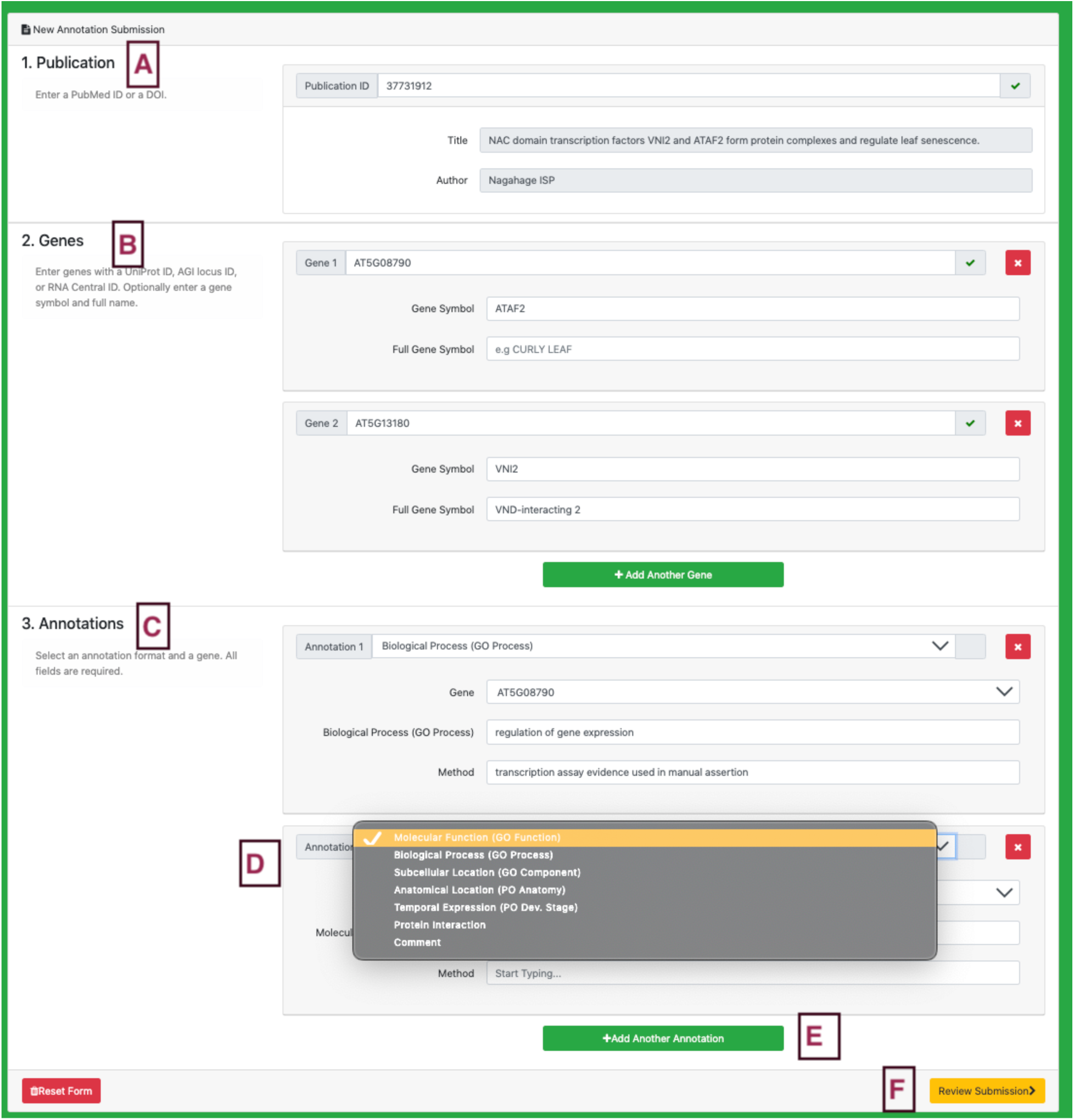
Screenshot of GOAT data submission interface after logging in via ORCiD. A) Users add DOI or PMID for the paper they are curating. B) Users enter locus identifiers and any gene names/symbols. Users can add more genes by clicking the ‘Add Another Gene’ button. C) Users must enter at least one annotation for at least one gene (specified in the above list). D) Users can add as many annotations as desired. E). They can choose different types of annotations from the drop down menu. The type of annotation determines the set of GO or PO terms available as well as the types of evidence (Method). Once all annotations are entered the user is prompted to review the submission (F) before submitting. Submissions are then reviewed by a TAIR curator before being imported into TAIR and integrated to the GO database on a quarterly basis.

GOAT was designed for flexibility and can be used to annotate ANY gene, from any organism, as long as it has an RNA central or UniProt ID. This provides the potential to fill in gaps for gene function in Arabidopsis or any plant species. Curating non-Arabidopsis gene function allows the capture of aspects of plant biology that either do not exist in Arabidopsis (such as nodulation or wood formation), or are more well described in other species. By extending curation of experimentally defined functions to other species we can narrow the knowledge gap and create a better representation of plant gene function across species.

## Website improvements

### Back end changes to improve database speed and stability

The Locus Detail pages are the most highly used pages at TAIR. For the time period spanning November 1, 2022 to October 31, 2023, our usage analytics show that locus pages were accessed 7,970,973 times. They consolidate all things gene function related: GO annotations, symbols and full names, summary, publications, alleles, germplasms, stock and clone information, RNA, protein and expression information, gene family and homolog data, as well as links out to external resources with complementary information about the genes. We were running into major issues with long page load times because of the substantial amount of data being retrieved from multiple tables in the TAIR Oracle database and then being aggregated. To solve this problem, we denormalized the Oracle data, and stored it in an Amazon Web Services Simple Storage Service (AWS S3) bucket as individual JSON files, which considerably sped up data processing and retrieval time and reduced computational costs.

AWS S3 is a scalable, durable and cost-effective object storage service provided by Amazon. It is a self-managed service, meaning that the organization using it (Phoenix Bioinformatics, in our case) doesn’t have to maintain physical servers. This leads to significant reductions in maintenance costs, as there’s no need to manage, upgrade, or replace server hardware. S3’s high scalability means that it can handle a growing volume of data and requests without compromising performance. AWS S3 is highly distributed, which means that data is redundantly stored across multiple data centers. This eliminates the risk of a single point of failure. One of the most crucial benefits of this migration to AWS S3 is the dramatic improvement in data retrieval speed. The data retrieval time for a typical Locus Detail page was reduced from one minute (using the previous technology) to an impressive 300 milliseconds when utilizing AWS S3. This is a substantial enhancement in user experience, making the Locus Detail pages much more responsive. In summary, by transitioning from traditional data storage and retrieval methods to denormalized data stored in AWS S3 buckets, we have not only achieved significant performance improvements but also reduced costs and improved the overall reliability and availability of the data.

We are currently working on a complete refactoring of the website using current technology and framework to replace the early 2000s-era software, with new modules deployed on the beta.arabidopsis.org website as they are completed.

### Moving to a better genome browser, JBrowse2

Since May 2020, JBrowse has been serving as TAIR’s primary genome browser. It is the source of the map images that we provide on both the Locus and Gene Detail pages. All new data tracks have been added exclusively to JBrowse since its introduction at TAIR. The Javascript-based JBrowse was first released in 2009 (Skinner *et al*. 2009) and is no longer being actively developed. To reduce the overall maintenance load associated with supporting four genome browsers (SeqViewer, GBrowse, JBrowse, and JBrowse2), support for the much older technology stack-powered SeqViewer and GBrowse will be discontinued. As more features and plugins are added to JBrowse2 and that platform becomes even more stable, we anticipate that we will shift exclusively to JBrowse2 as TAIR’s genome browser and sunset JBrowse support as well. User feedback to our announcement of sunsetting SeqViewer and GBrowse has highlighted several features of SeqViewer that are particularly valued by the community.

1. The ability to download gene sequences that have genome coordinates, UTRs, exons, introns and start and stop codons as well as intergenic regions marked. This can be done with the SeqLighter plugin in TAIR’s JBrowse and the feature has been requested from the JBrowse2 developers.
2. The ability to copy a DNA sequence from sequence viewer nucleotide view and paste to DNA editor applications like APE, keeping the lower case (intron and non-coding) and uppercase (exon) characters. This feature has been requested from the JBrowse2 developers.

JBrowse2 is a complete rewrite of JBrowse 1 with a similar user interface but a modern software architecture (Diesh *et al*. 2023). This more modern browser is under active development and maintenance, and features the capability for viewing genomic structural variants and evolutionary relationships among genes and genomes with syntenic visualizations. JBrowse2 is built on a contemporary tech stack and boasts optimized algorithms and a streamlined codebase, making it significantly faster than its predecessor. This means quicker load times, smoother navigation, and an overall enhanced user experience. The intuitive user interface makes the platform easier to navigate for seasoned users but also lowers the learning curve for newcomers. After testing JBrowse2 in beta mode for several months at TAIR, the tool is now available on the main website (jbrowse2.arabidopsis.org/index.html). Most of the data tracks that are available in the TAIR JBrowse are present in JBrowse2 (Figure 5). A few remain untransferred due to a current lack of the appropriate plugins for visualization and for that reason we will continue to maintain and update the original JBrowse.

**Figure 5:**
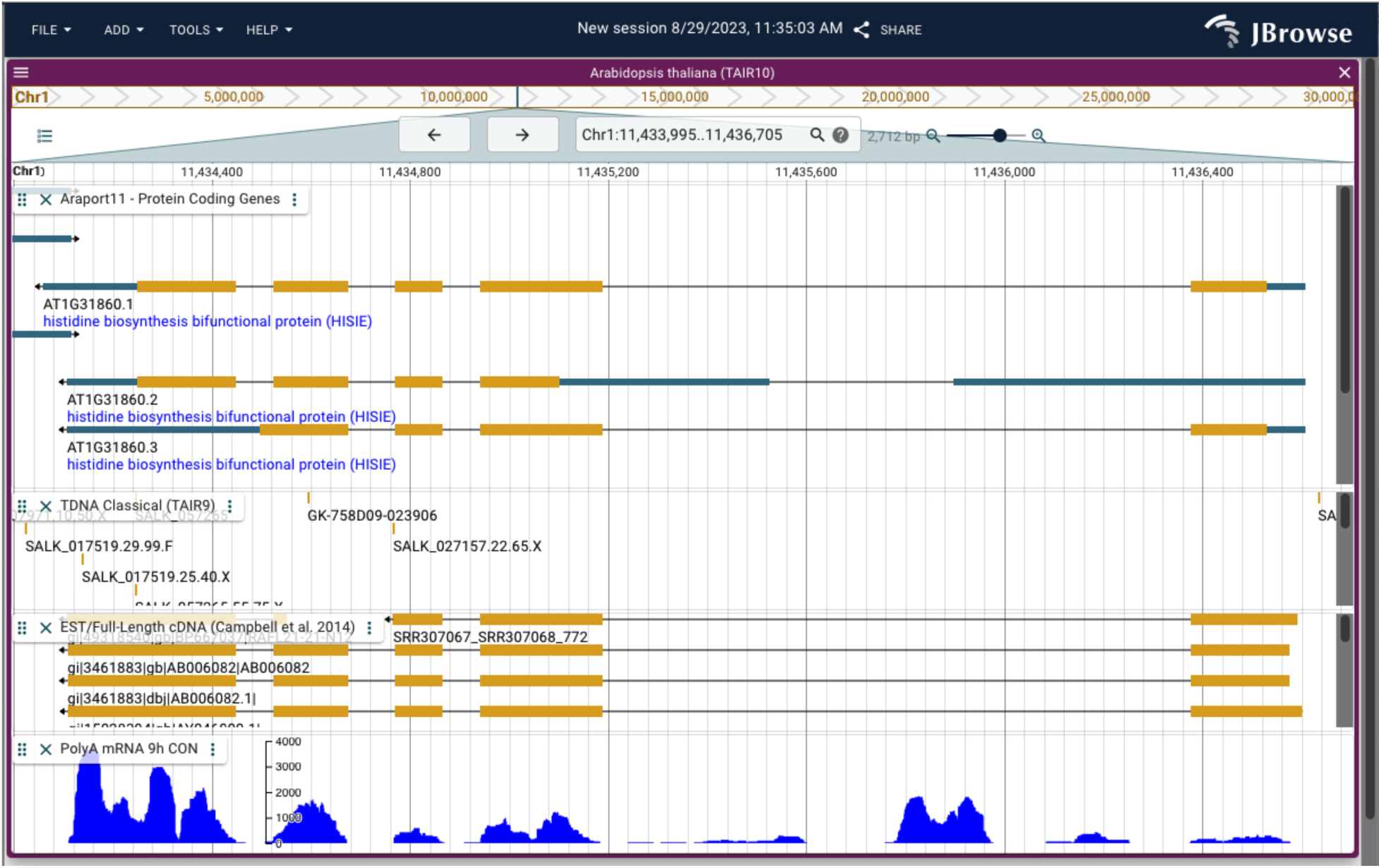
JBrowse2 interface showing the locus At1g31860, the structure of the three different gene models, locations of T-DNA insertions, supporting cDNAs and one some mRNA-seq expression data as a coverage track.

A new feature introduced in JBrowse2 enables visualization capability for syntenic datasets. This feature allows researchers to compare and contrast gene order and orientation across multiple genomes in a visually intuitive manner. By overlaying syntenic regions on the reference genome, JBrowse 2 provides a comprehensive view of genomic conservation, facilitating insights into evolutionary events such as gene duplications, inversions, and translocations. This integration of syntenic datasets into JBrowse 2 empowers scientists with a powerful tool for deciphering the complexities of genome evolution. In the initial release, we provide access to the *A. thaliana* and *A. lyrata* genomes for syntenic comparisons (Figure 6). Additional syntenic datasets for over 30 plant species, both monocot and dicot, will be made available as they are generated.

**Figure 6.**
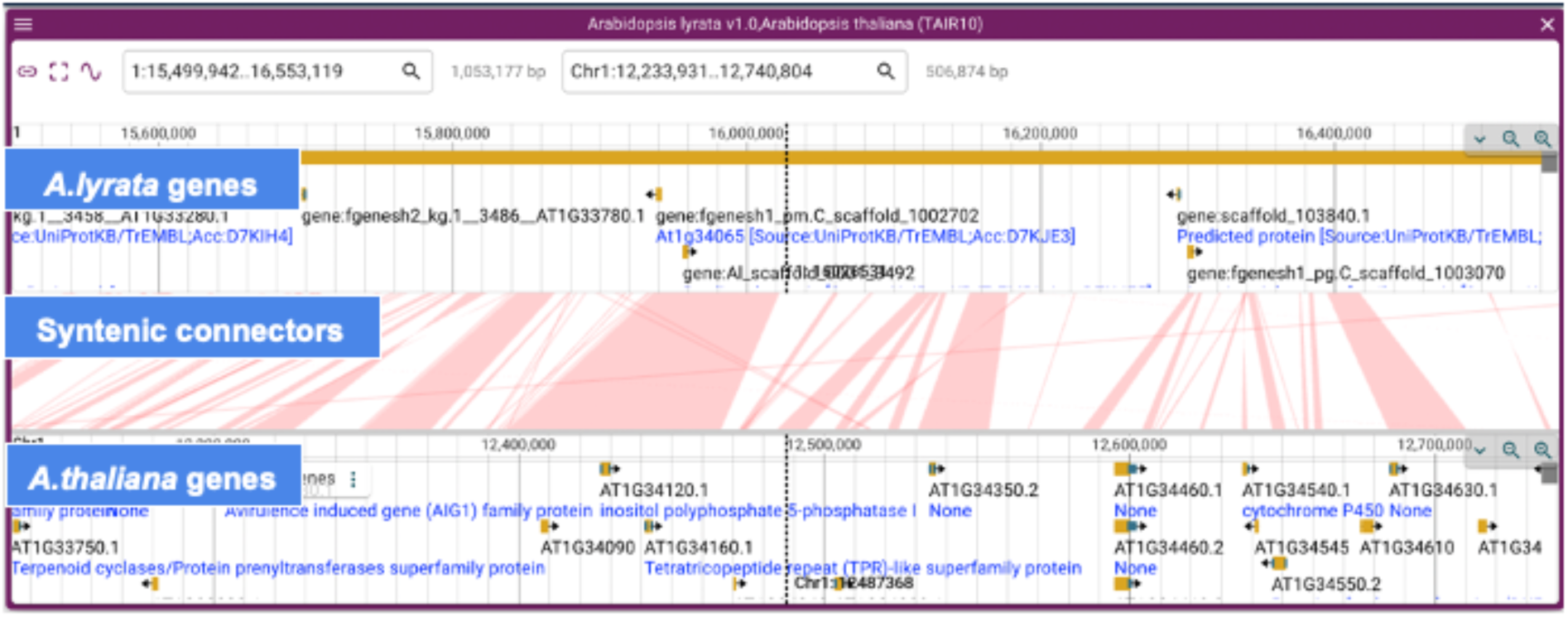
JBrowse 2 visualization of syntenic comparison between genomic regions of *A. thaliana* and *A. lyrata*. Individual tracks show *A. lyrata* protein coding genes (version 1.0), “Connectors” indicating syntenic regions between *A. thaliana* and *A. lyrata*, and *A. thaliana* protein coding genes (Araport11 release). Syntenic comparison was performed using MC-Scan and the output paf file was uploaded into JBrowse2 to create the above syntenic panel.

## TAIR as a community hub: Coordinating the reannotation of the genome

Since its inception, TAIR has functioned as a community hub for projects of broad interest, impact, and importance, such as maintaining project lists for the NSF 2010 project and the early days of the Multinational Arabidopsis Steering Committee (MASC). More recently, TAIR has played a key role in motivating, organizing and managing the latest reannotation of the reference genome based on the Col-0 ecotype.

### Brief history of genome releases

When the Arabidopsis thaliana genome was first sequenced and published in 2000, it marked the first complete plant genome available to the scientific community (Arabidopsis Genome Initiative 2000). After the initial annotation of the Col-0 genome, ten subsequent versions followed. The first four (TIGR2 through TIGR5) were done by what was then called The Institute for Genome Research (TIGR) (Haas *et al*. 2003, 2005).

The next five (TAIR6 through TAIR10) were funded by an NSF grant for the TAIR project (Swarbreck *et al*. 2008; Lamesch *et al*. 2012). Araport11 (Cheng *et al*. 2017), the most recent version that was released in June 2016 was funded by a grant to the J. Craig Ventner Institute (JCVI) for the Araport project. Over the years, additional experimental results like ESTs and RNAseq were incorporated into prediction pipelines with manual review following the automated predictions. The manual review process was not only necessary but essential in increasing the quality of each reannotation. The resulting products were incorporated into GenBank’s RefSeq section and from there were available to the broader bioinformatics and research community for use.

### Initiation of V12

With the advances in sequencing, assembly, and annotation technologies, it was glaringly apparent that an update to the Araport11 release was needed. Attempts to find directed funding for this effort were not successful and so another approach was needed. With the help of Nicholas Provart at the University of Toronto, Tanya Berardini, TAIR’s Director, convened a Zoom meeting of about 20 interested members from the Arabidopsis sequencing and genome assembly community in October 2022 to assess if there was broad support and community buy-in for a community resourced approach (<u>tinyurl.com/Athalianav12</u>). Scientific expertise would be provided by the community and project management, technical support and tool hosting would be provided by TAIR. Five phases of the annotation process were identified (Fig. 7): reference sequence assembly, automatic annotation, manual review, submission to the International Nucleotide Sequence Database Collaboration (INSDC), and dissemination/integration of the new reference into community tools and resources.

**Figure 7.**
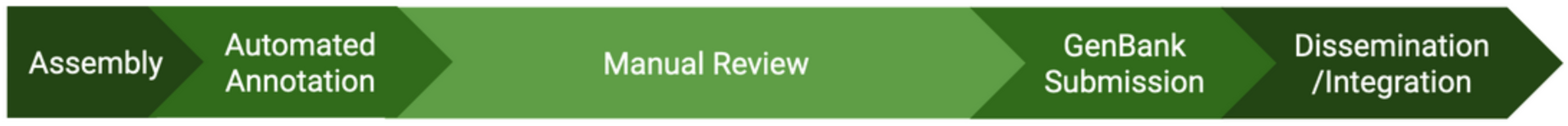
Phases of Genome Reannotation

The response was overwhelmingly in favor of proceeding with this plan. While the science of the genome reannotation will be reported in another publication (The Arabidopsis Col-CC Reannotation Team, in preparation), we wish to share some background on the organization of the effort as part of this update.

### Volunteer recruitment for all phases

Past annotation and reannotation efforts were done with dedicated funding to TIGR/TAIR/JCVI. Without dedicated funding for the 12th version, we were determined to make the most out of the Arabidopsis community’s expertise and goodwill to create a resource for the entire scientific community. Scientists across the globe were either recruited or came forward to contribute their skills in bioinformatics, annotation, assembly, automated annotation, manual review of genes, systematic reannotation of transposons and transposable elements, lncRNAs, rRNAs, and repeat elements. As much as possible, we tried to provide clear expectations and deadlines for completion of the assigned tasks. All contributors will be co-authors of the reannotation publication which will be submitted for review and publication after project completion. Contributions to the effort will be acknowledged using CRediT (Contributor Roles Taxonomy, https://credit.niso.org/), as they are for this publication.

### Apollo hosting

TAIR decided to use Apollo (Dunn *et al*. 2019) as our community curation tool for reviewing the results of the NCBI automated annotation pipeline. After initially setting up a very small test instance, the final Apollo server was an AWS EC2 instance with 16G memory, 2 vCPU and 1000 GB storage space, which provided not only enough storage space for all of the evidence tracks but also enough capacity to perform file manipulation and transformation. Most of the evidence tracks were provided to us in GFF format. Some needed to be transformed to bigwig format for more intuitive visualization and gene model validation.

### Manual Review Training

We were able to draw from the deep experience in the broader genome annotation community that has used Apollo for similar projects. Specifically, we reused and adapted teaching slides and guidelines from the maize community (shared by Marcela Tello Ruiz at Gramene), the Apollo developer group (shared by Monica Muñoz Torres, create while with Berkeley Bioinformatics Open-source Projects (BBOP)), and workflow management from the i5K project (shared by Monica Poelchau and Chris Childers at the US Department of Agriculture). We conducted six-1.5 hr training sessions over 5 weeks. Over 70 participants from 10 countries attended at least one of the sessions. Weekly Zoom office hours (one and a half hours a week, with two different times to accommodate global time zones) were established for ‘live’ feedback and troubleshooting and for the almost three months of manual review, at least one community member took advantage of the discussion time.

### Communication

With a globally distributed volunteer force, it was essential to have both synchronous and asynchronous communication channels. We established a central website for tracking progress and milestones (tinyurl.com/Athalianav12). Phoenix hosted a dedicated Slack channel for the manual review part of the project. Updates were shared by email, Slack, and TAIR’s X account. The combination of all of these venues were necessary to ensure that information was distributed in a timely fashion and reached the needed audiences. Regular updates kept the community motivated, involved, and informed.

In the process of organizing and executing the reannotation project we have identified useful tools, resources and strategies, as well as potential pitfalls to avoid. We plan to share the resources and lessons gleaned from this experience, to help other groups that may face similar challenges (i.e. lack of funding for genome annotation).

## Consolidating online community resources

Another way that TAIR serves as a community resource is by identifying and sharing useful data resources. The ‘Arabidopsis Community Resources Portal (https://conf.phoenixbioinformatics.org/display/COM/Resources) is a curated collection of databases, data sets and other digital resources of interest beyond TAIR for Arabidopsis researchers. The initial list was curated by members of the MASC Bioinformatics subgroup. Each entry is tagged with searchable keywords such as ‘gene_expression’, and ‘proteomics’ or entire list can be browsed via the page tree structure. We welcome suggestions and contributions from community members to add to this resource.

## Promoting FAIR standards as part of the AgBioData Consortium

TAIR also engages with the broader research community to promote better practices in data management and reuse. TAIR is a founding member of the AgBioData Consortium (www.agbiodata.org) and supports efforts to ensure that agricultural and related data are Findable, Accessible, Interoperable and Reusable (FAIR; (Wilkinson *et al*. 2016). Towards that end we participate in consortium-wide working groups (WGs) aimed at developing data and data management standards (Harper *et al*. 2018; Saha *et al*. 2022; Deng *et al*. 2023; Clarke *et al*. 2023). We have also drafted some guidelines for authors on how to make their Arabidopsis publications more FAIR (https://conf.phoenixbioinformatics.org/pages/viewpage.action?pageId=22807345;(Reiser *et al*. 2018)) and updated our list of recommendations on where to submit data including what data TAIR accepts and what it does not (https://www.arabidopsis.org/submit/index.jsp). We welcome feedback from the community.

## Perspective on future direction

Almost twenty five years after TAIR’s inception, the resource continues to grow and adapt to the changing needs of the community and the constantly shifting landscape of the technology that supports its online delivery. Changes in funding model aside, TAIR’s strength has always been and continues to be its deep connections with the scientific community that it serves. We will continue to nurture those ties and use them to guide TAIR’s expansion into new areas with the essential services that researchers and students have relied on for so many years.

### Long term sustainability

As a core resource for plant biologists, it is essential to have secure, long term funding. Since 2013 TAIR has been supported by community subscriptions, and has successfully transitioned away from episodic grant funding (Reiser *et al*. 2016). For the last decade,TAIR has been funded largely by over 225 academic institutional subscriptions (61% of TAIR’s total subscription revenue), a few national subscriptions (25%), and corporate subscriptions (10%) that cover full access to the resource for tens of thousands of scientists all over the world. There are also a couple hundred individual academic subscribers who contribute about 3% of TAIR’s total subscription revenue. Subscriptions have provided a stable source of funding that supports ongoing curation and some of the enhancements and improvements outlined here. Even with modest increases to cover inflation, the renewal rate for institutional subscriptions has been fairly stable (over 95%). We continue to offer the lowest rates that we can and provide free access for (1) teaching purposes (21 courses at 20 institutions in 2023 alone), (2) to US-based Historically Black Colleges and Universities, and (3) to countries classified as Low income economies by the World Bank. At this point, almost half of TAIR’s lifetime has been self-supported and we look forward to continuing to provide a valued, high quality resource to the community.

## Data Availability Statement

The website URL is www.arabidopsis.org. The GO annotation file available at doi.org/10.5281/zenodo.7843882 can be used to reconstruct Figure 1. The data files used for generating Figure 2 are available at FigShare. We will update the files on a regular basis with updates available through the TAIR website at https://www.arabidopsis.org/download/index-auto.jsp?dir=%2Fdownload_files%2FGenes%2FUnknown_Gene_Lists. Cumulative data files with information on gene function, publication links, germplasm and phenotype information, as well as GFF files with updated gene symbol and full name information are released every quarter (beginning of January, April, July, October). Subscriber Data Releases contain data updated within the last 12 months and are available at this URL to those with current subscriptions to TAIR: https://www.arabidopsis.org/download/index-auto.jsp?dir=/download_files/Subscriber_Data_Releases. Use of these files are governed by the Terms of Use, full text available here: http://www.arabidopsis.org/doc/about/tair_terms_of_use/417.

After a year, the Subscriber Data Releases are moved into the Public Data Releases folders at this URL: https://www.arabidopsis.org/download/index-auto.jsp?dir=/download_files/Public_Data_Releases

All files in the Public_Data_Releases folder are made available to the public under the CC-BY 4.0 license (https://creativecommons.org/licenses/by/4.0/).

## Database citation

If TAIR is either generally useful or essential in your research, please cite this publication (or any of the older TAIR publications from the reference list) whenever you publish your own work. Model organism databases provide a huge resource for the scientific community and their contributions are not recognized often enough in the published literature. Literature citation helps track not only TAIR’s but other MOD’s impact in a quantifiable manner. Such metrics are essential evidence in outreach efforts to funding agencies.

## Acknowledgements

The GOAT software was originally developed by the following Rochester Institute of Technology students: Austin Hartnett, John King, Ian Montgomery, Nick Peretti, Arron Reed, Ben Grawi, Gavin Nishizawa and integrated into Phoenix’s technology stack by Qian Li.

The authors thank the members of the plant biology research community for their continued support, feedback, and data contributions. This work was funded by national, academic, institutional, corporate and individual subscriptions to TAIR. Open access to this publication is funded by subscriptions.

## Conflict of interest section

None declared.

## Author contributions

Conceptualization: LR and TZB

Data curation: LR, EB, S. Subramaniam, and TZB

Formal Analysis: LR and TZB

Investigation: LR and TZB

Project administration: TZB

Software: XC, KK, S. Sawant, S. Subramaniam, and TP

Supervision: TZB and TP

Visualization: LR, S. Subramaniam, TZB

Writing – original draft: LR, TZB, TP, S. Sawant, and S. Subramaniam

Writing – review & editing: all authors

## References

Arabidopsis Genome Initiative, 2000 Analysis of the genome sequence of the flowering plant Arabidopsis thaliana. Nature 408: 796–815. 10.1038/35048692

Arnaboldi V., D. Raciti, K. Van Auken, J. N. Chan, H.-M. Müller, et al., 2020 Text mining meets community curation: a newly designed curation platform to improve author experience and participation at WormBase. Database 2020. 10.1093/database/baaa006

Ashburner M., C. A. Ball, J. A. Blake, D. Botstein, H. Butler, et al., 2000 Gene ontology: tool for the unification of biology. The Gene Ontology Consortium. Nat. Genet. 25: 25–29. 10.1038/75556

Berardini T. Z., D. Li, R. Muller, R. Chetty, L. Ploetz, et al., 2012 Assessment of community-submitted ontology annotations from a novel database-journal partnership. Database 2012. 10.1093/database/bas030

Berardini T. Z., L. Reiser, D. Li, Y. Mezheritsky, R. Muller, et al., 2015 The Arabidopsis Information Resource: Making and Mining the “Gold Standard” Annotated Reference Plant Genome. Genesis 53: 474–485. 10.1002/dvg.22877

Cheng C.-Y., V. Krishnakumar, A. P. Chan, F. Thibaud-Nissen, S. Schobel, et al., 2017 Araport11: a complete reannotation of the Arabidopsis thaliana reference genome. Plant J. 89: 789–804. 10.1111/tpj.13415

Clarke J. L., L. D. Cooper, M. F. Poelchau, T. Z. Berardini, J. Elser, et al., 2023 Data sharing and ontology use among agricultural genetics, genomics, and breeding databases and resources of the AgBioData Consortium. arXiv [cs.DB].

Deng C. H., S. Naithani, S. Kumari, I. Cobo-Simon, E. H. Quezada-Rodriguez, et al., 2023 Agricultural sciences in the big data era: Genotype and phenotype data standardization, utilization and integration. Preprints.

Diesh C., G. J. Stevens, P. Xie, T. De Jesus Martinez, E. A. Hershberg, et al., 2023 JBrowse 2: a modular genome browser with views of synteny and structural variation. Genome Biol. 24:74. 10.1186/s13059-023-02914-z

Dunn N. A., D. R. Unni, C. Diesh, M. Munoz-Torres, N. L. Harris, et al., 2019 Apollo: Democratizing genome annotation. PLoS Comput. Biol. 15: e1006790. 10.1371/journal.pcbi.1006790

Garcia-Hernandez M., T. Z. Berardini, G. Chen, D. Crist, A. Doyle, et al., 2002 TAIR: a resource for integrated Arabidopsis data. Funct. Integr. Genomics 2: 239–253. 10.1007/s10142-002-0077-z

Gaudet P., M. S. Livstone, S. E. Lewis, and P. D. Thomas, 2011 Phylogenetic-based propagation of functional annotations within the Gene Ontology consortium. Brief. Bioinform. 12: 449–462. 10.1093/bib/bbr042

Gene Ontology Consortium, S. A. Aleksander, J. Balhoff, S. Carbon, J. M. Cherry, et al., 2023 The Gene Ontology knowledgebase in 2023. Genetics 224: iyad031. 10.1093/genetics/iyad031

Haas B. J., A. L. Delcher, S. M. Mount, J. R. Wortman, R. K. Smith Jr, et al., 2003 Improving the Arabidopsis genome annotation using maximal transcript alignment assemblies. Nucleic Acids Res. 31: 5654–5666. 10.1093/nar/gkg770

Haas B. J., J. R. Wortman, C. M. Ronning, L. I. Hannick, R. K. Smith Jr, et al., 2005 Complete reannotation of the Arabidopsis genome: methods, tools, protocols and the final release. BMC Biol. 3: 7. 10.1186/1741-7007-3-7

Harper L., J. Campbell, E. K. S. Cannon, S. Jung, M. Poelchau, et al., 2018 AgBioData consortium recommendations for sustainable genomics and genetics databases for agriculture. Database 2018: bay088. 10.1093/database/bay088

Huala E., A. W. Dickerman, M. Garcia-Hernandez, D. Weems, L. Reiser, et al., 2001 The Arabidopsis Information Resource (TAIR): a comprehensive database and web-based information retrieval, analysis, and visualization system for a model plant. Nucleic Acids Res. 29: 102–105.

Jacobson M., A. E. Sedeño-Cortés, and P. Pavlidis, 2018 Monitoring changes in the Gene Ontology and their impact on genomic data analysis. Gigascience 7. 10.1093/gigascience/giy103

Jones P., D. Binns, H.-Y. Chang, M. Fraser, W. Li, et al., 2014 InterProScan 5: genome-scale protein function classification. Bioinformatics 30: 1236–1240.

Kishore R., V. Arnaboldi, C. E. Van Slyke, J. Chan, R. S. Nash, et al., 2020 Automated generation of gene summaries at the Alliance of Genome Resources. Database 2020. 10.1093/database/baaa037

Lamesch P., T. Z. Berardini, D. Li, D. Swarbreck, C. Wilks, et al., 2012 The Arabidopsis Information Resource (TAIR): improved gene annotation and new tools. Nucleic Acids Res. 40: D1202–1210. 10.1093/nar/gkr1090

Larkin A., S. J. Marygold, G. Antonazzo, H. Attrill, G. Dos Santos, et al., 2021 FlyBase: updates to the Drosophila melanogaster knowledge base. Nucleic Acids Res. 49: D899–D907. 10.1093/nar/gkaa1026

Li D., T. Z. Berardini, R. J. Muller, and E. Huala, 2012 Building an efficient curation workflow for the Arabidopsis literature corpus. Database 2012. 10.1093/database/bas047

Müller H.-M., K. M. Van Auken, Y. Li, and P. W. Sternberg, 2018 Textpresso Central: a customizable platform for searching, text mining, viewing, and curating biomedical literature. BMC Bioinformatics 19: 94. 10.1186/s12859-018-2103-8

Nadendla S., R. Jackson, J. Munro, F. Quaglia, B. Mészáros, et al., 2022 ECO: the Evidence and Conclusion Ontology, an update for 2022. Nucleic Acids Res. 50: D1515–D1521. 10.1093/nar/gkab1025

Paniagua C., A. Bilkova, P. Jackson, S. Dabravolski, W. Riber, et al., 2017 Dirigent proteins in plants: modulating cell wall metabolism during abiotic and biotic stress exposure. J. Exp. Bot. 68: 3287–3301. 10.1093/jxb/erx141

Reiser L., T. Z. Berardini, D. Li, R. Muller, E. M. Strait, et al., 2016 Sustainable funding for biocuration: The Arabidopsis Information Resource (TAIR) as a case study of a subscription-based funding model. Database 2016. 10.1093/database/baw018

Reiser L., L. Harper, M. Freeling, B. Han, and S. Luan, 2018 FAIR: A Call to Make Published Data More Findable, Accessible, Interoperable, and Reusable. Mol. Plant 11: 1105–1108. 10.1016/j.molp.2018.07.005

Reiser L., S. Subramaniam, P. Zhang, and T. Berardini, 2022 Using the Arabidopsis Information Resource (TAIR) to Find Information About Arabidopsis Genes. Curr Protoc 2: e574. 10.1002/cpz1.574

Rocha J. J., S. A. Jayaram, T. J. Stevens, N. Muschalik, R. D. Shah, et al., 2023 Functional unknomics: Systematic screening of conserved genes of unknown function. PLoS Biol. 21: e3002222. 10.1371/journal.pbio.3002222

Rutherford K. M., M. A. Harris, A. Lock, S. G. Oliver, and V. Wood, 2014 Canto: an online tool for community literature curation. Bioinformatics 30: 1791–1792. 10.1093/bioinformatics/btu103

Saha S., S. Cain, E. K. S. Cannon, N. Dunn, A. Farmer, et al., 2022 Recommendations for extending the GFF3 specification for improved interoperability of genomic data. arXiv [q-bio.OT].

Skinner M. E., A. V. Uzilov, L. D. Stein, C. J. Mungall, and I. H. Holmes, 2009 JBrowse: a next-generation genome browser. Genome Res. 19: 1630–1638. 10.1101/gr.094607.109

Swaminathan K., K. Peterson, and T. Jack, 2008 The plant B3 superfamily. Trends Plant Sci. 13: 647–655. 10.1016/j.tplants.2008.09.006

Swarbreck D., C. Wilks, P. Lamesch, T. Z. Berardini, M. Garcia-Hernandez, et al., 2008 The Arabidopsis Information Resource (TAIR): gene structure and function annotation. Nucleic Acids Res. 36: D1009–14. 10.1093/nar/gkm965

The Gene Ontology Consortium, 2019 The Gene Ontology Resource: 20 years and still GOing strong. Nucleic Acids Res. 47: D330–D338. 10.1093/nar/gky1055

The UniProt Consortium, A. Bateman, M.-J. Martin, S. Orchard, M. Magrane, et al., 2023 UniProt: the Universal Protein Knowledgebase in 2023. Nucleic Acids Res. 51: D523– D531. 10.1093/nar/gkac1052

Wilkinson M. D., M. Dumontier, I. J. J. Aalbersberg, G. Appleton, M. Axton, et al., 2016 The FAIR Guiding Principles for scientific data management and stewardship. Sci Data 3: 160018.

Xue B., and S. Y. Rhee, 2023 Status of genome function annotation in model organisms and crops. Plant Direct 7: e499. 10.1002/pld3.499

Zhang P., T. Z. Berardini, D. Ebert, Q. Li, H. Mi, et al., 2020 PhyloGenes: An online phylogenetics and functional genomics resource for plant gene function inference. Plant Direct 4: e00293. 10.1002/pld3.293

